# Maternofetal transfer of human NMDAR-antibodies leads to cortical network defect in adolescence

**DOI:** 10.1101/2023.11.30.569370

**Authors:** Saleh Altahini, Jan Doering, Joseph Kuchling, Hendrik Backhaus, Jakob Kreye, Roberta Guimaraes-Backhaus, Carsten Finke, Harald Prüss, Albrecht Stroh

## Abstract

Anti-N-methyl-D-aspartate-receptor IgG-antibodies (NMDAR-Ab) can be detected in up to 1% of the healthy population. Because maternal IgG can cross the placental barrier during pregnancy, these antibodies may influence fetal neurodevelopment. Using a mouse model of maternofetally transferred human NMDAR-Ab, we asked whether *in utero* exposure impacts functional brain dynamics later in life. We conducted two-photon calcium imaging and fMRI in adolescent mice, to evaluate both local and global neuronal network integrity. At the microcircuit scale, NMDAR-Ab exposed offspring exhibited lower spontaneous activity, reduced firing variability, and reduced orientation selectivity upon visual stimulation. Globally, fMRI revealed a selective bilateral functional hypoconnectivity of the entorhinal-hippocampal pathway while overall functional connectivity is unaffected. Together, our data indicates that maternal-fetal transfer of NMDAR-Ab leads to a lasting impairment in sensory processing and memory-routing circuits, revealing a vulnerability of the developing fetal brain to maternal autoantibodies.

## Introduction

Maintaining stable local and brain-wide functional states is essential for mental health. These mental-health-associated functional dynamics are established very early in life. Already *in utero*, particularly the balance between excitatory and inhibitory activity starts forming (Ben-Ari 2002, Xu, Broadbelt et al. 2011). Neurodevelopmental trajectories are highly sensitive to maternal immune cues, and transient perturbations during pregnancy can lead to long-lasting structural and functional changes in the offspring across their lifespan. Severe maternal immune activation during active infections constitutes a risk factor for neuropsychiatric disorders (Estes and McAllister 2016, Yong, Zhao et al. 2025). Yet non-infectious autoimmune pathways may pose an additional significant risk to the developing fetal brain.

The interactions between the brain and the immune system are bidirectional. The activation of the immune system during active infections leads to sickness behavior (Dantzer 2004, Filiano, Gadani et al. 2015). As the cytokines released by immune cells interact with neural networks, they promote adaptive behaviors that help combat the disease, such as reduced water and food consumption, decreased physical activity, and a diminished inclination towards social interaction (Dantzer, O’Connor et al. 2008). However, in autoimmune conditions, the immune system produces autoantibodies that can directly target and modify neural circuits. This has sparked a debate whether traditional psychiatric conditions, such as major depressive disorder and schizophrenia, are primarily disorders of neural circuitry or if they are driven by chronic systemic inflammation and autoimmune pathways (Dantzer, O’Connor et al. 2008, Carvalho, Torre et al. 2013, Khandaker, Cousins et al. 2015, Miller and Raison 2016, Howes, Bukala et al. 2024).

A key example of a neuropsychiatric disorder caused directly by autoantibodies is anti-NMDA receptor encephalitis (NMDARE) (Dalmau, Tuzun et al. 2007). IgG antibodies that specifically bind to the N-methyl-D-aspartate receptors (NMDAR) are the cause of NMDARE. Although NMDARE is considered to be a rare disease, anti-NMDA receptor antibodies (NMDAR-Ab) can also be detected in the blood of up to 1% of healthy subjects (Dahm, Ott et al. 2014, Lang and Pruss 2017). Consequently, a considerable number of women may carry these anti-neuronal antibodies and are at risk of transferring them to the fetus during pregnancy. This exposure may not lead to NMDARE, however, the exposure to the antibodies leads to the internalization of the receptors and thereby impact synaptic transmission. In early stages of pregnancy, the fetal blood-brain-barrier (BBB) is still permeable (Kumar, Jain et al. 2010) and the IgG placental transfer has already started (Palmeira, Quinello et al. 2012). This allows for the interaction of antibodies with fetal brain tissue potentially leading to altered brain development *in utero*.

NMDA receptors play a key role in long-term potentiation (LTP) and thus in modulating synaptic plasticity and controlling memory functions (Li and Tsien 2009, Luscher and Malenka 2012). In humans, NMDAR dysfunction has been linked to dementia and neurodegeneration (Choi, Koh et al. 1988, Zhou, Hollern et al. 2013, Wang and Reddy 2017) as well as psychotic experience (Coyle 2012, Sterzer, Adams et al. 2018). It has been well established that NMDAR-Abs cause an internalization of the receptors (Kreye, Wenke et al. 2016, Masdeu, Dalmau et al. 2016). In the maternofetal transfer mouse model of NMDAR-Ab, exposure reduced the cortical density of NMDAR and led to behavioral changes in the offspring that can persist until adulthood (Jurek, Chayka et al. 2019, Garcia-Serra, Radosevic et al. 2021). Resting state-fMRI showed significantly reduced functional connectivity of hippocampal subregions even at the age of 10 months (Kuchling, Jurek et al. 2024).

Here, we ask whether NMDAR-Ab exposure due to maternofetal transfer leads to a persistent change in local cortical and brain-wide networks in the adolescent mouse. To investigate these developmental defects across different brain scales, we focused on both local sensory microcircuits and brain-wide functional networks. For that, we used the murine model of *in utero* exposure to human recombinant NMDAR-Ab (Jurek, Chayka et al. 2019). We employed two-photon calcium imaging in the awake mouse to probe local network integrity with single-neuron resolution, affording to resolve subtle network state changes (Arnoux, Willam et al. 2018, Ellwardt, Pramanik et al. 2018, Rosales Jubal, Schwalm et al. 2021). We assessed a local microcircuit in layer II/III of the primary visual cortex. Recording in primary sensory cortical areas enables capturing both, spontaneous, task-free activity, mirroring the functional microarchitecture of a given network, and the precision of sensory coding upon sensory stimulation. Brain-wide functional integrity was assessed by functional Magnetic Resonance Imaging (fMRI). By combining single-neuron two-photon calcium imaging in V1 with resting-state fMRI, we determined whether *in utero* NMDAR-Ab exposure impacts both local sensory processing and global network connectivity. We identified a distinct, maladaptive brain state in NMDAR-Ab exposed mice at local and brain-wide scale in the adolescent offspring, long after the *in utero* exposure.

## Material and Methods

### Animals

Animal experiments were carried out in accordance with institutional animal welfare guidelines and were approved by the local ethics committee for Animal Welfare (Landesuntersuchungsamt Rhineland-Palatinate, LaGeSO Berlin). In the treatment group, pregnant C57BL/6 mice were injected intraperitoneally at gestational days E13 and E17, at each timepoint with 240 μg of human monoclonal IgG1 antibody #003-102. Controls were exposed to a maternofetal transfer of human monoclonal IgG1 antibody that is non-reactive to brain tissue (control clone: #mGO53). The sample size for this study was determined by the litter size of pregnant mothers and the probability of a high SNR functional imaging experiment per mouse. All animals were kept on a 12h day/night cycle. Food and water were given *ad libitum*.

### Virus preparation, injection, and surgery

A viral solution containing AAV1.CamKII.GCaMP6f.WPRE.SV40 (virus titer: 0,367*10^12^/ml) was used. Virus injection and chronic window implantation were performed for all animals age-matched on postnatal day 24 and according to our STAR Protocol (Guimaraes Backhaus, Fu et al. 2021). During the entire procedure, body temperature and respiration rate were continuously monitored. To start, the mouse was first weighed and then placed in an anesthesia induction chamber (4% vol/vol isoflurane/O^2^ induction, 1-2% sustained). To minimize the pain, a diluted solution of buprenorphine (0.1 mg/kg body weight) was injected subcutaneously before transferring the animal to the stereotaxic frame. The head was then shaved, and a local anesthetic (2% xylocaine) was applied to the scalp. To protect the eyes, a 5% dexpanthenol eye cream was applied. An incision was then made at midline with a scalpel blade and extended to a circular cut with surgical scissors to clear the skull. The edges of the scalp were sealed with surgical glue. The periosteum was scraped off with the scalpel blade and the skull was thoroughly cleaned with sponges and saline solution. To improve the adhesion of the head holder to the skull, small, grid pattern grooves were made with the scalpel blade and then with a dental drill. A 2.5 mm craniotomy above V1 was drilled 2.7 mm posterior of Bregma and 2.3 mm lateral of the midline. Approx. 1 µL of viral solution were then injected using a fine micropipette prepared specially for intracortical injections at cortical depth of ∼ 150-250 µm targeting layer II/III of V1. After that, a 3.0 mm glass cover was placed over the craniotomy and sealed with surgical glue. The head holder was then attached to the skull using UV-glue. After the surgery, all animals received *ad libitum* access to buprenorphine in drinking water and were observed closely for the first 24 hours then checked for the following two days.

### Two-photon imaging experiments

Awake GCaMP6f-imaging was performed on P41 using a custom-made two-photon microscope (TrimScope II by LaVision Biotec) equipped with Ti:Sapphire laser (Chameleon II from Coherent) and a resonant scanner at a wavelength of 920 nm. The imaging plane was usually between 250 +/- 100 µm below the cortical surface. A Zeiss W-Plan-Apochromatic 20 x DIC VIS-IR (NA=1.0) water-immersion objective was used to achieve a field of view of 465×465 µm. Images were sampled at a rate of 30.8 Hz and a size of 512×512 pixels. The frame trigger signal of the microscope was sampled using a data acquisition interface (Cambridge Electronic Design Limited) at 2000 Hz and was later used to sync frame timings with visual stimulation.

### Visual stimulation

For visual stimulation, 10 trials of static and drifting gratings were generated by a custom written Python script and projected onto Phenosys 270° virtual reality system (6 thin-film transistor (TFT) monitors arranged in a hexagonal shape). Each trial consisted of 5 s uniform grey screen followed by a randomized series of 8 gratings with different orientations, each was presented for 10 s (5 s static + 5 s drifting). The orientations ranged from 0° to 315° with a step size of 45°. The order of presentation was logged to a text file and was later synchronized to the functional calcium imaging by aligning the microscopes frame trigger signal to the time points of the presentations.

### Two-photon data analysis

Analysis of functional calcium imaging datasets was performed using CaImAn and a custom-built analysis pipeline. To determine optimal parameters for CaImAn processing (including patch size, patch overlap, and the number of components per patch), a subset of recordings was randomly selected and manually inspected using a custom script. Because all imaging data were acquired using identical microscope settings and objective, the optimized parameters were kept constant across all recordings to ensure consistency. Raw imaging data underwent rigid and non-rigid motion correction, source extraction (segmentation), and ΔF/F trace extraction via CaImAn. The resulting spatial footprints, denoised traces, and raw ΔF/F traces were exported and stored in an HDF5 file structure. Peak detection was carried out via a semi-automated pipeline using a custom GUI application. Regions of interest (ROIs) were initially filtered to exclude low-quality components, automatically discarding any ROI with a CaImAn SNR below 3 or a convolutional neural network classifier prediction (cnn_preds) below 0.15. For the remaining valid ROIs, the raw ΔF/F traces were temporally smoothed using a 1D Gaussian filter (σ=1). Dynamic event thresholds were determined using a localized noise estimator calculated from the lower-half distribution of the smoothed ΔF/F signal. Specifically, baseline noise was defined by scaling the distance between the absolute minimum and the median of the positive-valued trace. The prominence and height thresholds for peak detection were then set to a minimum of ten times this estimated noise floor. Automated peak detection was executed on normalized, denoised traces using the scipy.signal.find_peaks function. To prevent the false splitting of complex, multi-peaked transients, a minimum temporal separation distance was enforced between consecutive peaks, defined dynamically as the product of the estimated calcium indicator decay time.

Following automated detection, a manual quality control step was used to visually review the overall reliability of each segmented ROI. This stage focused on a global assessment to exclude ROIs displaying obvious motion artifacts or false positives, and to re-include true somatic components that marginally failed the automated filters, without modifying or re-evaluating individual transient profiles. To standardize downstream computational analyses and minimize the influence of lingering baseline noise or indicator decay kinetics, we constructed a simplified binarized activity matrix for each ROI. For every confirmed transient, the segment spanning from the detected onset to its corresponding peak was preserved, while the uninformative decay phase was ignored. Rather than a strictly binary (0 or 1) value, active frames within this onset-to-peak window were assigned a continuous value scaled linearly between 0 and 1, relative to the maximum detected peak amplitude across all events recorded for that specific ROI.

Burstiness was determined using the Coefficient of Variation:

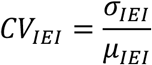

σ*_IEI_*: The standard deviation of the inter-event intervals (measuring temporal variability/dispersion). μ*_IEI_*: The mean of the inter-event intervals (measuring the average time gap between events). A *CV_IEI_* > 1 denotes irregular, burst-dominated firing, whereas *CV_IEI_* ≈ 1 represents random Poisson-like activity. *CV_IEI_* < 1 indicates more regular firing than a Poisson process (e.g., pacemaker-like activity).

The Circular Variance (CV) (Ringach, Shapley et al. 2002) was calculated as the variance of the cell’s response to all orientations:

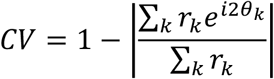

Where *r_k_* is the cells response to a given orientation, *r_k_* is the angle of the grating in radians. The oblique effect was assessed by comparing the proportion of tuned cells between cardinal (0°, 90°, 180°, 270°) and oblique (45°, 135°, 225°, 315°) orientations.

### MRI data acquisition and quality control

Anesthesia was achieved using 1.5-2% isoflurane in a 70:30 nitrous oxide:oxygen mixture. Before start of the rsMRI scan, isoflurane levels were reduced to 1.2 (+/- 0.3) % until animals breathed regularly at higher rate of 160 (+/-25)/min. This process took 5-7 min and body temperature and respiration rate were monitored with MRI compatible equipment (Small Animal Instruments Inc., Stony Brook, NY). 2D echo-planar imaging (EPI) resting-state functional MR (rs-fMRI) images were acquired (repetition time (TR) = 1000 ms; echo time (TE) = 13 ms; flip angle (FA) = 50°; 300 repetitions; 16 axial slices with slice thickness = 0.75 mm; field of view (FOV) = 19.2 x 12.0 mm²; image matrix = 128 x 80; number of averages = 1) on a 7 T MR scanner (Bruker Biospec, Ettlingen, Germany) and a transmit/receive mouse cryoprobe (Bruker). Experimenters were blinded to the condition of the animals.

### rs-fMRI data processing, denoising and analysis

Processing of rs-fMRI data was modified according to an established rs-fMRI analysis protocol (Zerbi, Grandjean et al. 2015). Rs-fMRI datasets were skull-stripped using linear affine registration of an in-house mouse atlas template based on C57BL/6 control mice previously studied at the same scanner to individual mouse rs-fMRI images with FSL FLIRT (Jenkinson, Bannister et al. 2002). Each skull-stripped 4D dataset was then fed into FSL MELODIC (Beckmann and Smith 2004) to perform within-subject spatial independent component analysis (ICA) with automatic dimensionality estimation, using a skull-stripped EPI image of a deliberately chosen subject as anatomical template. Within-subject ICA was performed after removal of the first 10 out of 300 time points of every subject and included in-plane smoothing with a 1.0 x 1.0 mm kernel. Subsequent denoising of single-subject fMRI independent components was performed in accordance with previously described protocols (Zerbi, Grandjean et al. 2015, Griffanti, Douaud et al. 2017) by manual identification of signal and noise based on the threshold spatial maps, the temporal power spectrum, and the time course by one rater (J.K.) who was blinded to the clinical phenotype of mice. For functional connectivity analysis, the brain was segmented according to the anatomical atlas published by Dorr et al (Dorr, Lerch et al. 2008). Eleven regions corresponding to the cerebral ventricles and major white matter tracts were excluded from analysis to restrict the functional connectome to gray-matter structures, resulting in 103 ROIs. Regional BOLD timeseries were extracted, and pairwise functional connectivity, network complexity, graph topology, and dynamic brain states were calculated as described in the unified analysis section below. The resting-state fMRI cohort consisted of 15 NMDAR-Ab exposed offspring and 15 controls.

### Network Complexity, Graph Topology, and State Dynamics Analysis

To evaluate the functional relationships between network nodes, functional connectivity (FC) was computed for both data modalities. For two-photon calcium imaging, FC was defined as the cosine similarity between the binarized somatic activity vectors of all neuron pairs. For rs-fMRI, FC was computed as the Pearson correlation coefficient between the BOLD timeseries of all 103 Dorr atlas regions.

Network complexity was evaluated using Principal Component Analysis (PCA) and Non-Negative Matrix Factorization (NMF). The population dimensionality was defined as the minimum number of principal components (PCs) required to explain at least 80% of the total network variance. Spatial co-active ensembles were extracted by fitting a 5-component NMF model. To ensure non-negativity for NMF, raw calcium traces were clipped at zero, while fMRI BOLD timeseries were baseline-shifted by subtracting each region’s minimum value. The spatial organization of these ensembles was quantified using the Hoyer sparsity index of the spatial weight matrix (H).

Functional networks were constructed by thresholding the individual FC matrices (cosine similarity for two-photon, Pearson’s r for fMRI) across five proportional densities: 5%, 10%, 15%, 20%, and 25%. Graph metrics were calculated at each density and averaged. We assessed the average clustering coefficient (C); characteristic path length (L) and the small-worldness index (S). Hub nodes were defined as nodes whose degree exceeded the network mean by more than two standard deviations.

Temporal network dynamics were modeled by clustering the population activity vectors (all cells or regions at each frame) into *_K_* = 4 discrete brain states using K-Means clustering. Continuous denoised calcium traces (C traces) were used for two-photon state transitions, and denoised BOLD timeseries were used for rs-fMRI. State transition dynamics were characterized by calculating the average state dwell time (the consecutive duration spent in a state before transitioning, measured in seconds or repetition times) and the transition entropy (in bits), which quantifies the mathematical randomness of state transitions. Transition entropy was calculated as the average Shannon entropy of the transition probability distributions across all *_K_* = 4 states:

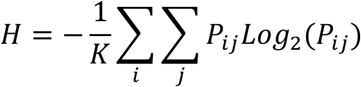

where *P_ij_* is the probability of transitioning from state *_i_* to state *_j_*. Measured in bits, a transition entropy of 0 denotes a completely deterministic network, whereas a maximum value of 2 bits represents a completely random, uniform transition structure.

### Statistical tests

Statistical analysis was done with GraphPad Prism. All datasets were first tested for normal distribution, in cases where a normal distribution was rejected, a non-parametric test was done. All statistical tests and exact p-values are noted under the corresponding figures. Box-and-whisker plots indicate the median value (center line), the 25–75th percentiles (box) and the 10–90th percentiles (whiskers). All other graphs represent mean ± standard error of the mean (S.E.M.) unless otherwise stated. The significance level was set at p < 0.05.

## Data availability

The data supporting the findings of this study are available upon request. Scripts used for the analysis can be accessed on our GitHub repository https://github.com/StrohLab.

## Results

### NMDAR antibody exposure leads to a decrease in the spontaneous activity in visual cortical microcircuits six weeks after NMDA exposure *in utero*

To examine the long-term effects of maternofetal transfer of anti-NMDAR antibodies on neuronal microcircuit dynamics of mouse primary visual cortex (V1), we injected pregnant C57BL/6 mice with 240 μg of NMDAR-Ab solution on gestational day E13 and again on day E17 (**Fig. 1A**). On the postnatal day 24, we stereotactically injected a viral solution encoding for GCaMP6f, targeting layer II/III of V1, alongside with the implantation of a metal head holder and chronic window for awake imaging. GCaMP6f-expression was controlled by the CaMKII-promoter, targeting only excitatory neurons. At adolescence (P41) we conducted two-photon imaging in layer II/III of V1 of awake behaving mice (**Fig. 1B**). We did not find significant differences in the number of active cells per recording (**Fig. 1C**) and the signal to noise ratio (SNR) (**Fig. 1D**) comparing mice exposed to NMDAR-Ab and control animals. This indicates that morphologically, the network is intact, devoid of apparent cell loss.

**Fig. 1:**
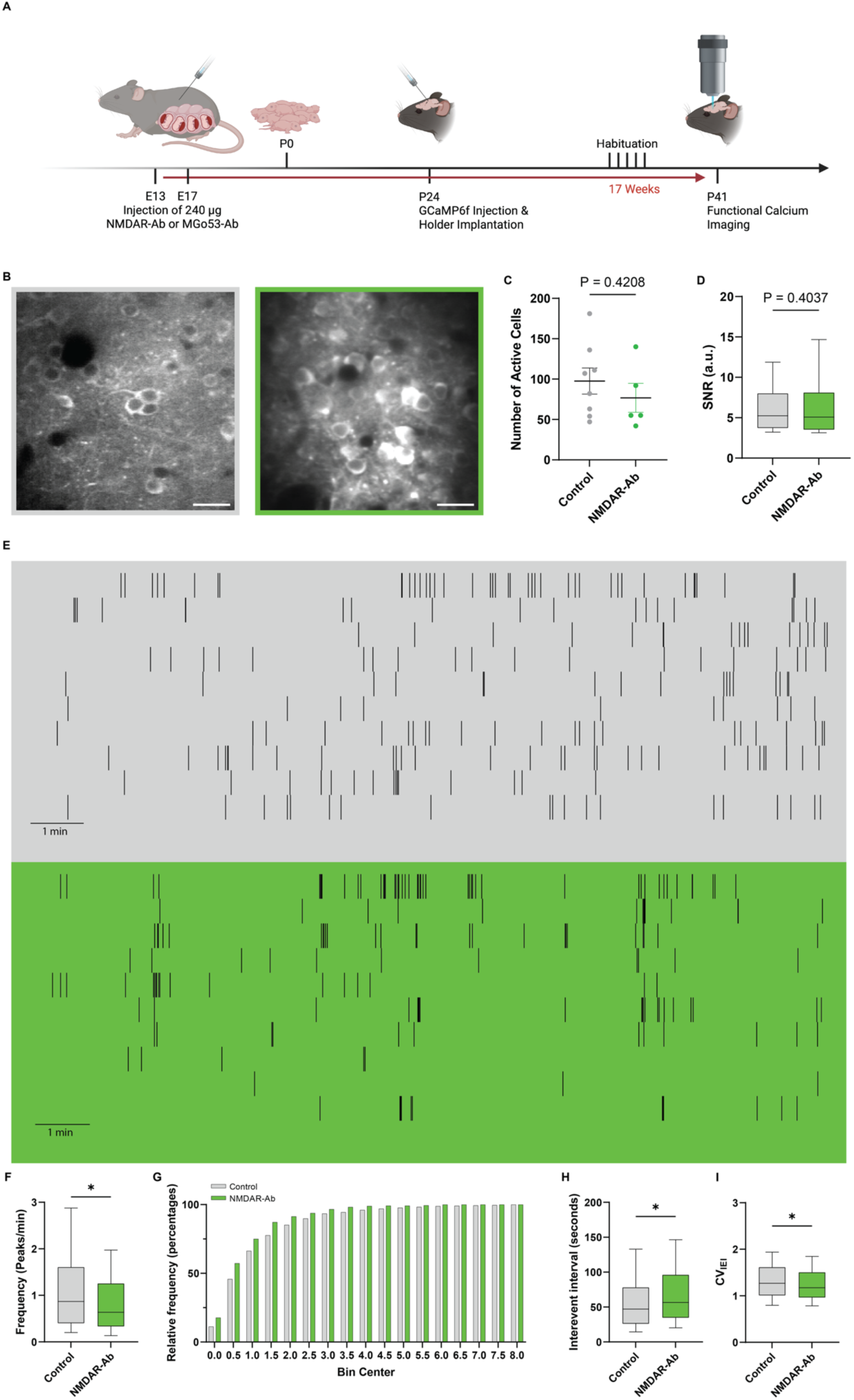
Maternal-fetal transfer of NMDAR autoantibodies reduces spontaneous microcircuit firing rate and irregular dynamics in offspring. (**A)** The timeline of the experiments: For maternofetal transfer of antibodies, pregnant C57BL/6 mice were injected with 240 µg of NMDAR-Ab solution on gestational day E13 and E17. On postnatal day 24, the injection of GCaMP6f viral solution and implantation of a chronic window were performed. 3 weeks after the surgery, the mice were habituated to the experimental setup. Imaging was conducted on P41. (**B)** Two-photon imaging fields from Control (left) and NMDAR-Ab (right) (Magnified view from the center of 20x field of view. Scale bar 50 µm). (**C)** No significant difference in total number of active cells per field of view, (Unpaired t-test, p=0.4208). (**D)** No significant difference in the SNR of active neurons of control and NMDAR-Ab animals. (Mann Whitney test, p=0.4037) (**E)** Representative binary activity matrices (raster plots) showing the temporal dynamics of somatic calcium events over time for 10 randomly selected active neurons in a representative Control (**top**) and NMDAR-Ab exposed (**bottom**) microcircuit. Each row represents a single neuron, and ticks indicate calcium transient peak. **(F)** Firing frequency of active cells under spontaneous conditions, showing a significant decrease in NMDAR-Ab exposed animals (median: 0.6354 vs 0.8695 peaks/min in control; Mann Whitney test, p<0.0001). **(G)** Cumulative frequency distribution of the spontaneous activity frequencies from panel F. **(H)** Interevent interval (IEI) distributions showing a significant shift towards longer intervals in NMDAR-Ab offspring (median: 56.57 vs 47.06 s in control; Mann-Whitney test p=0.0002). **(I)** Firing irregularity of active cells quantified by the coefficient of variation of interevent intervals (*CV_IEI_*), demonstrating more rigid dynamics in NMDAR-Ab exposed animals (1.173 vs 1.268 in Control; Mann-Whitney test, p=0.0084). Sample sizes: Control group: n=781 cells from regions from 8 unique regions/7 animals; NMDAR-Ab group: n=384 cells from 5 unique regions/4 animals.

We probed the functional architecture of ongoing, spontaneous activity, in adolescent mice that had been exposed to NMDAR-Ab approx. six weeks earlier. Our analysis yielded a binary activity matrix of the recorded cells (**Fig. 1E**), similar to a spiking matrix of electrophysiological recordings. In the NMDAR-Ab group, we found a reduced mean frequency of spontaneous Ca^2+^ transients (**Fig. 1F**). Splitting up the activity levels into activity bins in a cumulative probability graph, a rather homogenous shift towards lower activity bins became apparent in NMDAR-Ab exposed animals (**Fig. 1G**). This indicates a rather microcircuit-wide reduction in activity, in both high and low active neurons, paralleled by an increase in the inter-event intervals (IEI) (**Fig. 1H**). NMDAR-Ab exposed networks showed a reduction in the variability of firing, i.e. a more homogenous firing pattern (**Fig. 1I**). Both the overall reduction in activity, and the divergent temporal pattern of activity suggests a stable new setpoint of the functional microarchitecture in cortical circuits upon *in utero* NMDAR-Ab exposure, impacting only distinct features of network function.

### V1 excitatory neurons of NMDAR-Ab exposed animals exhibit a lower visual tuning

We next exposed the animals to a visual stimulation paradigm consisting of a randomized sequence of 5s drifting and static gratings (**Fig. 2B**) in a 270° virtual reality setup (**Fig. 2A**). Particularly layer II/III of visual cortex is characterized by highly tuned neurons, i.e., neurons responding to only one or few orientations of the gratings. While in both groups, highly tuned neurons could be observed (**Fig. 2C**), the representation of visual afferents was altered in NMDAR-Ab exposed mice (**Fig. 2D, E**). Exposed offspring exhibited a significantly higher Circular Variance (CV), corresponding to lower overall orientation tuning capability relative to controls. We now asked whether the reduction of the degree of tuning is a mere consequence of the overall lower activity state. We therefore shuffled the timing of the Ca^2+^ transient, while preserving the overall number of calcium transients. The shuffling led to a drastic decrease of tuning in both groups, indicating an active, adaptive to maladaptive network state shift in the NMDAR-Ab exposed group.

**Fig. 2:**
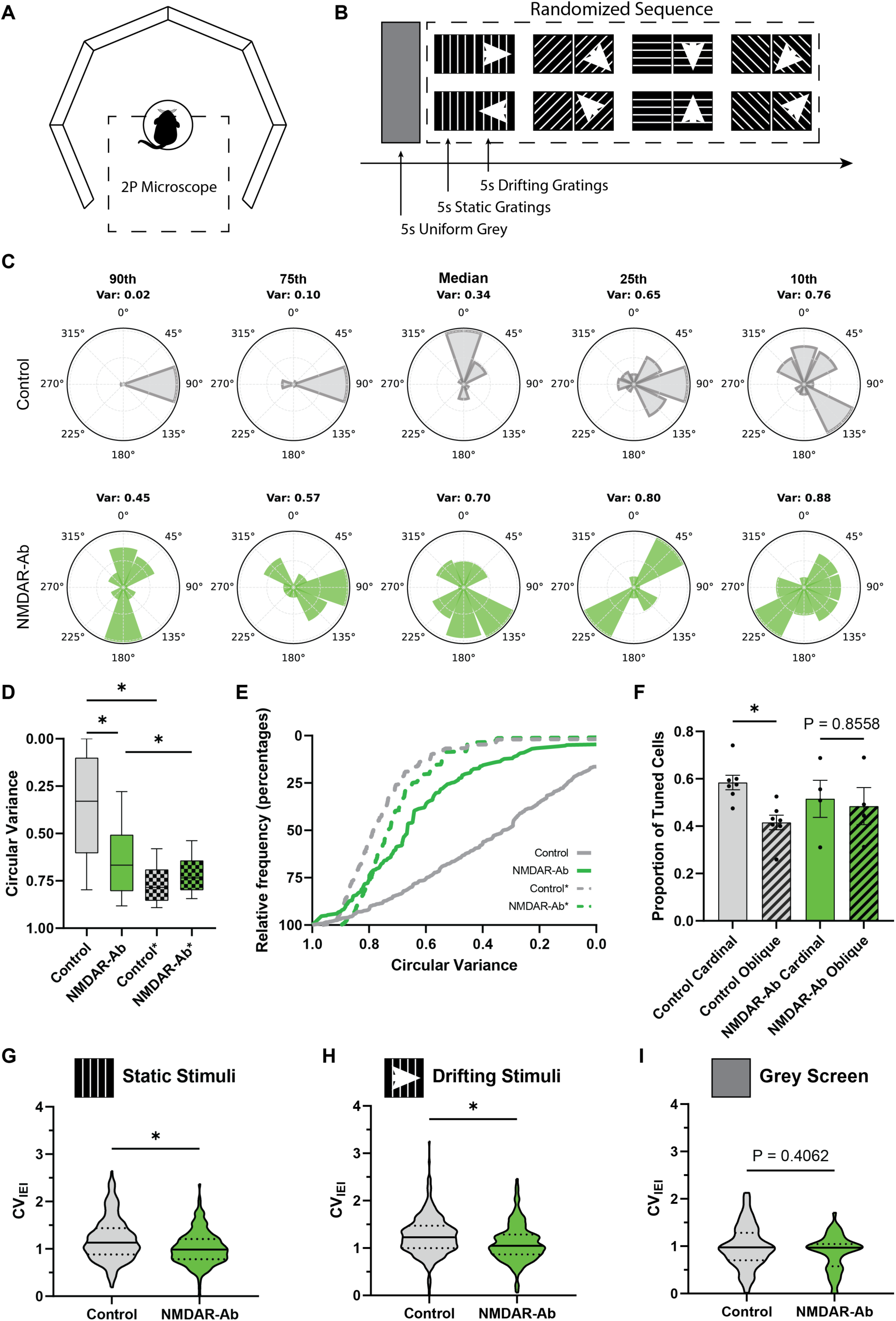
In utero NMDAR-Ab exposure reduces orientation selectivity in cortical microcircuits. **(A)** Illustration of the virtual reality setup. The mouse is head mounted under the microscope on a Styrofoam trackball and can move freely in all directions. A 270° surrounding monitor system is placed around the mouse for visual stimulation. **(B)** Illustration of the visual stimulation paradigm. Each trial starts with 5 seconds of uniform grey screen followed by a randomized sequence of gratings at different orientations, where each orientation is displayed for 5 seconds statically and then 5 seconds in drifting motion. **(C)** Exemplary orientation tuning plots of five cells from control (top) and NMDAR-Ab (bottom) mice sampled at the 90th (maximum), 75th, 50th (median), 25th, and 10th (minimum) percentiles of orientation selectivity. **(D)** Higher circular variance (CV) reveals a lower orientation tuning of V1 neurons in NMDAR-Ab compared to control and temporally shuffled controls (*=randomized, Kruskal-Wallis test, Control vs NMDAR-Ab p<0.0001, Control vs. Control* p<0.0001, NMDAR-Ab vs NMDAR-Ab* p=0.0237). **(E)** Cumulative distribution of the circular variance. (F) Proportion of active cells showing a preferred tuning for cardinal (0°, 90°, 180°, 270°) and oblique (45°, 135°, 225°, 315°). Control group shows a significant preference for cardinal orientations over oblique orientations (paired t-test, p=0.0330). NMDAR-Ab group shows no significant difference in tuning proportions between cardinal and oblique orientations (paired t-test, p=0.856). **(G-I)** Firing irregularity of active cells quantified by the coefficient of variation of interevent intervals (*CV_IEI_*) during distinct visual stimuli. **(G)** NMDAR-Ab exposed microcircuits exhibit significantly lower burst firing compared to controls upon presentation of static gratings (median: 0.986 vs 1.131 in Control; Mann-Whitney test, p<0.0001). **(H)** Burst firing is significantly reduced in the NMDAR-Ab cohort upon presentation of drifting gratings (median: 1.050 vs 1.230 in Control; Mann-Whitney test, p<0.0001). **(I)** Baseline firing irregularity remains stable and is not significantly different between groups during grey screen periods (median: 0.968 vs 0.974 in Control; Mann-Whitney test, p=0.406). Sample sizes: Control group: n= 698 cells from regions from 7 unique regions/7 animals); NMDAR-Ab group: n=303 cells from 4 unique regions/3 animals.

This visual processing deficit was further characterized by a breakdown of normal sensory bias architecture. Control animals displayed a classic, expected preference for cardinal orientations (0°, 90°, 180°, 270°) over oblique orientations (45°, 135°, 225°, 315°), illustrating standard cortical network asymmetry (**Fig. 2F**). This bias is known as the oblique effect in network science (Appelle 1972). This physiological effect was entirely abolished in the NMDAR-Ab exposed group. What is more, NMDAR-Ab exposed animals exhibited a reduced firing variability (**Fig. 2G–I**).

### Functional connectivity architecture of local circuits is largely unaffected

We characterized the functional connectivity architecture as Pearson’s correlation coefficient between neuronal pairs (**Fig. 3A, B**). While visually, NMDAR-Ab exposed animals seem to exhibit a more homogenous connectivity, no statistically significant quantitative shift could be determined. We now asked, whether NMDAR-Ab exposure alters the underlying state space of network dynamics. In this context, the number of principal components (PCs) required to explain 80% of the total variance of the activity matrix serves as a direct proxy for network dimensionality. A high number of PCs indicates a complex network with highly diverse, independent neural configurations, whereas a lower number implies a simpler and more redundant network. This analysis revealed that the functional dimensionality of the network state-space is not significantly affected by NMDAR-Ab exposure (**Fig. 3C**).

**Fig. 3:**
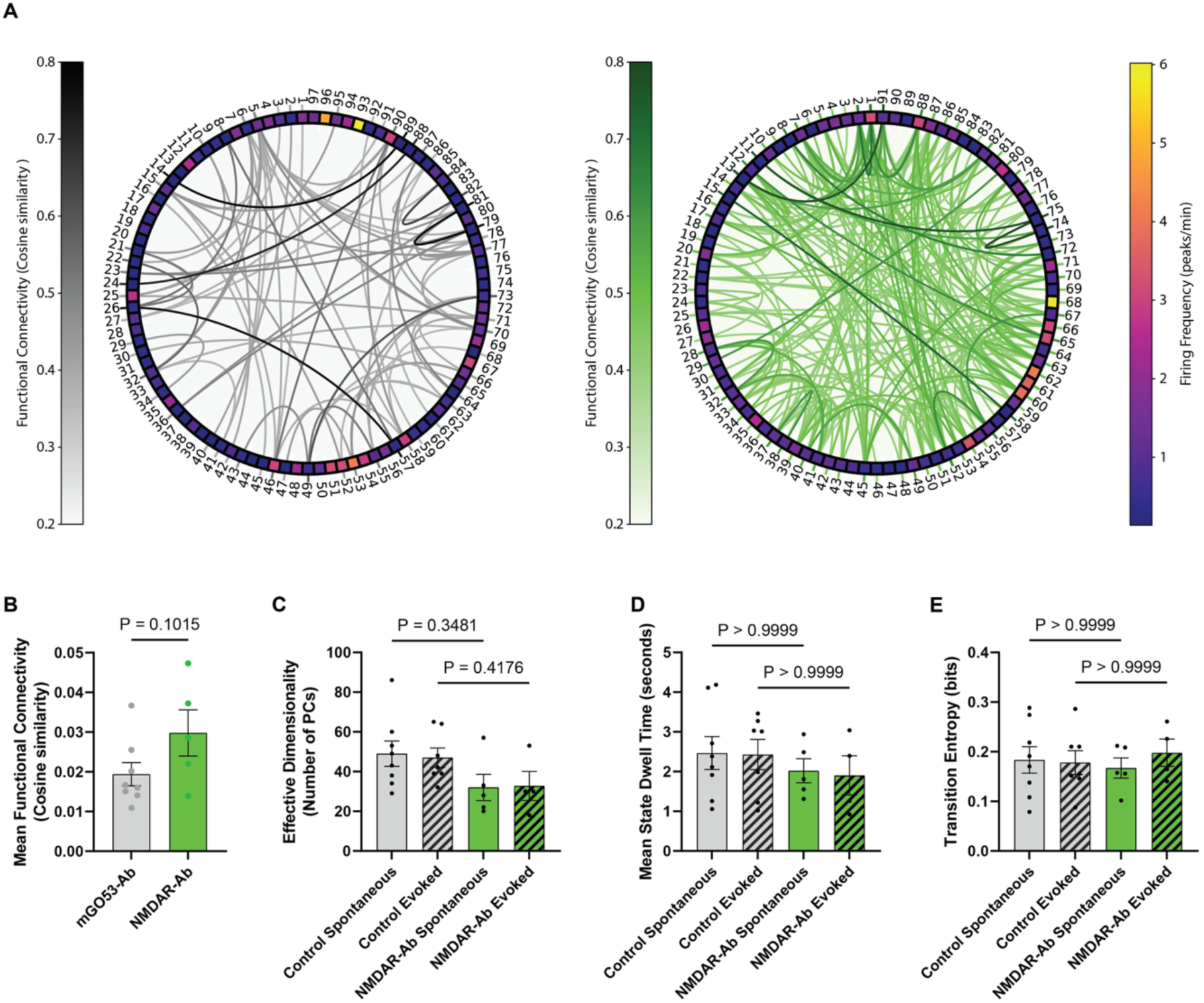
Local microcircuit network complexity, graph topology, and state transitions remain stable under NMDAR-Ab exposure. **(A)** Representative correlation matrices showing functional connectivity layout in Control (**left**) and NMDAR-Ab (**right**) offspring. **(B)** Mean Functional Connectivity (Pearson’s r) under spontaneous conditions, showing a trend towards increased average correlation in the NMDAR-Ab group (mean: 0.0298 vs. 0.0194 in Control; unpaired t-test, p=0.102). **(C)** PCA Network Dimensionality (number of PCs explaining >= 80% variance), showing non-significant trend-level decreases in NMDAR-Ab offspring under both spontaneous (mean: 32.00 vs. 49.00 in Control; Holm-Šídák test, p=0.348) and visually evoked activity (mean: 32.75 vs. 47.00 in Control; Holm-Šídák test, p=0.418). **(D)** Mean State Dwell Times show no significant differences between groups under spontaneous (Dunn’s test, adjusted p > 0.999) or visually evoked activity (Dunn’s test, p>0.999). **(E)** State Transition Entropy shows stable levels of transition randomness under both spontaneous (p>0.999) and visually evoked states (p>0.999, Dunn’s test). Sample sizes are the same as in figures 1 and 2.

We next assessed the shifts between functional states over time. We applied a k-means clustering algorithm to partition the high-dimensional population activity into discrete, recurrent network states. To determine the optimal number of states, we applied the elbow method across all datasets and experimental groups, which pointed to k = 4 states. While a few datasets showed an optimal cluster size of k = 5, we explicitly limited the analysis to 4 states across all recordings to maintain consistency and allow for direct statistical comparisons. Using these 4 defined states, we tracked the mean state dwell times, i.e. how long the network remains in a specific state before switching. We also assessed the total state transition entropy. Neither the mean state dwell times (**Fig. 3D**) nor the total state transition entropy was significantly altered by the *in utero* exposure (**Fig. 3E**). Subsequent graph theoretical analyses revealed no significant differences between groups as well (**Fig. S1**).

### NMDAR-Ab exposure does not lead to an apparent change in brain-wide network dynamics

Using fMRI, we observed the presence of typical resting state networks, such as the default mode network (**Fig. 4A**). Next, we conducted a parcellation approach, subdividing the brain in 103 regions based on anatomical brain atlas for MRI analysis (**Fig. 4B**). Applying our analysis pipeline on the resulting BOLD activity traces yielded no significant changes in the dimensionality of brain-wide networks (**Fig. 4C**). We then used k-means clustering to partition the network into discrete activity states, again limiting k to 4 states based on previous testing. The results showed no significant changes in both the mean state dwell times (**Fig. 4D**) nor in the state transition entropy (**Fig. 4E**). Graph theoretical analysis of the networks did not show significant differences across experimental groups (**Fig. S2**).

**Fig. 4:**
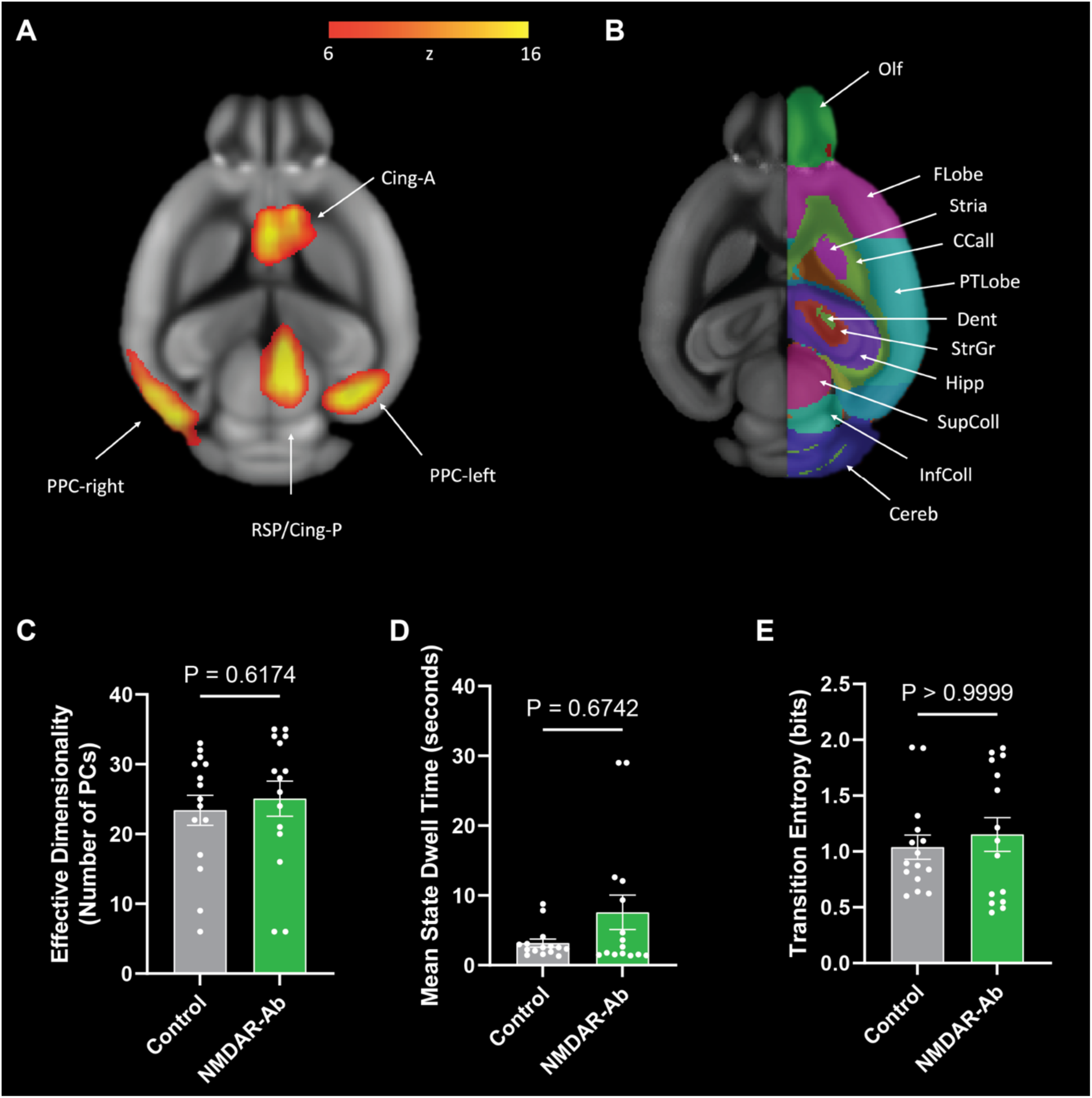
In utero NMDAR-Ab exposure preserves macro-scale complexity, graph topology, and brain states in rs-fMRI. **(A)** Activation pattern corresponding to the default mode network in an exemplary animal **(B)** fMRI parcellation into anatomical regions using a brain atlas. **(C)** PCA Network Dimensionality (number of PCs required to explain >= 80% BOLD variance) is stable between groups (mean: 25.07 vs. 23.40 in Control; unpaired t-test, p=0.617). **(D)** Mean Brain State Dwell Times show no significant differences between groups (median: 2.661 vs. 2.544 in Control; Mann-Whitney test, p=0.674). **(E)** State Transition Entropy, representing transition randomness, is stable between groups (median: 1.097 vs. 0.8940 in Control; Mann-Whitney test, p>0.999). Sample sizes: n=15 animals per group.

### Brain-wide networks in NMDAR-Ab exposed animals are characterized by pathway specific functional disconnections

We then quantified the functional connectivity between brain regions by Pearson’s correlation. We averaged the functional connection strength across all animals in each group to create network graphs (**Fig. 5A, B**). The mean functional connectivity across all regions was not significantly altered in the NMDAR-Ab group (**Fig. 5C**). We binned the functional connections into hemispheric subcategories (intra-hemispheric, inter-hemispheric, Left-Left and Right-Right). Comparing Control and NMDAR-Ab animals revealed no significant differences of the connection strength in those subcategories (**Fig. 5D**). However, a fine-grained, pathway-by-pathway analysis of individual functional connections revealed localized network deficits (**Fig. 5E**). Specifically, we identified selective, highly significant drops in functional connectivity centered on the entorhinal cortex bilaterally. This hypoconnectivity in NMDAR-Ab exposed offspring was prominent across entorhinal-hippocampal and subcortical projections (**Fig. 5F**).

**Fig. 5:**
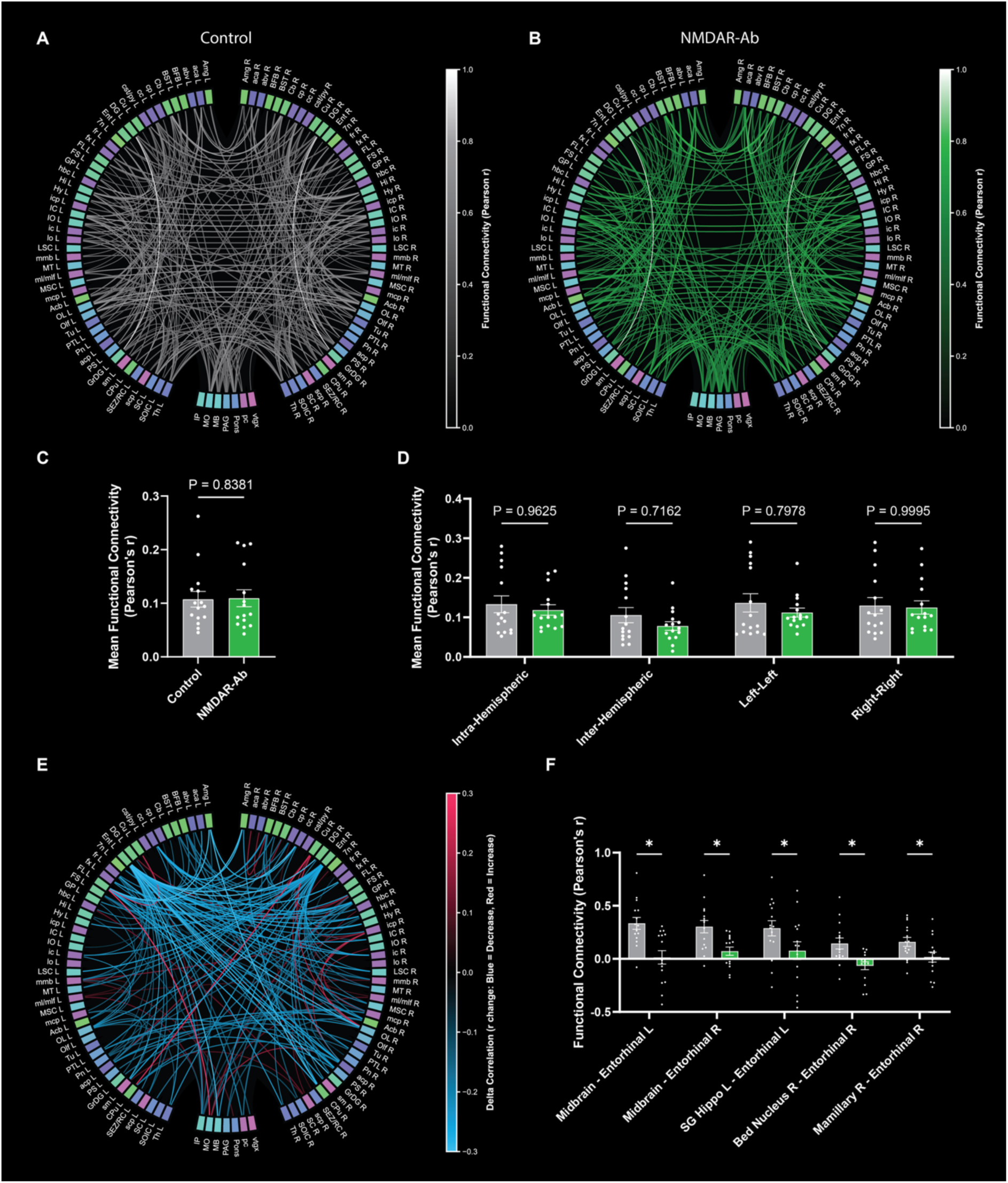
Resting-state BOLD fMRI reveals selective entorhinal-hippocampal functional disconnection. (A–B) Circular functional connectivity networks thresholded at r >= 0.45 for Control **(A)**, NMDAR-Ab exposed **(B)**. **(C)** Mean functional connectivity (average Pearson correlation coefficient) is similar between groups (median: 0.07951 vs. 0.09422 in Control; Mann-Whitney test, p=0.838). **(D)** Hemispheric functional connectivity comparing Control and NMDAR-Ab groups across intra-hemispheric (mean: 0.1186 vs. 0.1332 in Control; Šídák’s test, adjusted p=0.963), inter-hemispheric (mean: 0.0781 vs. 0.1057 in Control; Šídák’s test, adjusted p=0.716), Left-Left (mean: 0.1122 vs. 0.1366 in Control; Šídák’s test, adjusted p=0.798), and Right-Right connections (mean: 0.1250 vs. 0.1297 in Control; Šídák’s test, adjusted p=1.000). **(E)** Circular functional connectivity graph showing significant differences (p < 0.05) between groups **(F)** Pairwise functional connectivity of the 5 key entorhinal-hippocampal connections, showing a highly significant bilateral drop in NMDAR-Ab offspring: Midbrain - L. Entorhinal (p=0.00069), Midbrain - R. Entorhinal (p=0.00412), SG Hippo L - L. Entorhinal (p=0.00163), Bed Nucleus R - R. Entorhinal (p=0.00350), and Mamillary R - R. Entorhinal (p=0.00361; Welch’s t-tests). Sample sizes: n=15 animals per group

## Discussion

Our study reveals that a maternofetal transfer of NMDAR-Ab leads to multiple lasting cortical and brain-wide network defects in adolescent offspring. These deficits include aberrant spontaneous network dynamics as well as reduced precision in the encoding of sensory afferents. First, we report that *in utero* NMDAR-Ab exposure leads to a reduced cortical activity level, accompanied by a reduction of firing variability. Second, we found that V1 excitatory neurons exhibit a significant reduction in visual tuning and the loss of normal preference for cardinal orientations. Third, while the overall brain-wide functional connectivity remains largely intact, resting-state fMRI reveals a selective, bilateral functional disconnection of the entorhinal-hippocampal pathway. Together, these results demonstrate that transient prenatal exposure to maternal NMDAR-Ab leads to long-lasting brain state maladaptation in both local sensory processing and memory-routing networks.

### Early, in utero, exposure leads to a long-term maladaptive network state

One particular aspect of our study represents the large developmental interval between the administration of the NMDAR-Ab at E13 and E17, roughly comparable to the 5^th^ and 15^th^ gestational week in humans (Otis and Brent 1954), and the assessments of the network state in P41, roughly corresponding to adolescences (Cottam, Ofori et al. 2024). Our data shows that even a short transient exposure to NMDAR-Ab can lead to long lasting defects. Early, *in utero*, exposure to plasticity-mediating factors seems to lead to a long-term maladaptive brain state. The reduction of variability in the spontaneous activity constitutes a deviation from the typical divers and information-heavy functional architecture of cortical networks.

We chose primary visual cortex to examine local circuit effects as it shows highly organized response properties, such as orientation selectivity, that depend heavily on developmental NMDAR activity (Ramoa, Mower et al. 2001). After the transient exposure to NMDAR-Ab, V1 excitatory neurons displayed a significant reduction in orientation tuning selectivity, indicated by higher circular variance. The maternal antibodies likely disrupt this developmental refinement in utero, leaving the adult microcircuits with less selective, noisier representations of visual inputs.

Particularly the diminished oblique effect in the NMDAR-Ab exposed animals stands out. This effect refers to the bias towards a visual stimulus with a horizontal or vertical orientation (Appelle 1972) and is observed in many species, including humans and mice. It is believed that the oblique effect increases the network’s efficiency in encoding visual representations, as cardinal orientations are more frequently occurring in nature (Henderson and Serences 2021). It therefore seems likely, that the visual cortical circuits of NMDAR-Ab-exposed animals deviate substantially from the evolutionary preserved encoding principles of health-associated network function.

### Distinct local functional microarchitecture characterized by low, less variable firing

The local functional calcium imaging data revealed a circuit-wide transition to a state of low spontaneous activity in NMDAR-Ab exposed mice. This is in sharp contrast to findings in early states of neurological disorders, such as Alzheimer’s Disease, Multiple Sclerosis and Huntington’s Disease. Here, we and others reported a shift towards hyperactivity early in disease development (Arnoux, Willam et al. 2018, Ellwardt, Pramanik et al. 2018, Burgold, Schulz-Trieglaff et al. 2019, Rosales Jubal, Schwalm et al. 2021, Sosulina, Mittag et al. 2021, Wei, Xue et al. 2021). This dysregulation was brought about by a new functional group of hyperactive cells, which impacted local connectivity measures. Here, local connectivity remained largely unperturbed. This indicates that the developmental impact of the antibodies is rather homogenous across the local network, and leads to a new, quiet, stable network state.

### Scale-specific maladaptations: Brain-wide networks exhibit a regionally confined shift in connectivity

Overall brain-wide functional connectivity was unaffected by *in utero* NMDAR-Ab exposure as determined by resting-state fMRI. Yet, we identified a highly selective, bilateral hypoconnectivity within the entorhinal-hippocampal pathway. The entorhinal cortex acts as the primary interface between the neocortex and the hippocampus, routing spatial, sensory, and temporal information crucial for memory consolidation and navigation (Sasaki, Leutgeb et al. 2015, Hernández-Frausto and Vivar 2024). NMDARs are highly expressed in these circuits and are vital for synaptic plasticity, such as long-term potentiation, and the development of spatial grid cell networks (Gil, Ancau et al. 2018, Beesley, Sullenberger et al. 2019). Our findings suggest that transient exposure to maternal NMDAR-Abs during gestation selectively impairs the wiring or functional maturation of these entorhinal-hippocampal projections. This pathway-specific hypoconnectivity provides a clear structural and functional correlate for the long-term memory impairments previously observed in similar animal models (Garcia-Serra, Radosevic et al. 2021) and highlights a potentially higher vulnerability of the developing temporal lobe to maternal NMDAR-Ab exposure.

### Translational Implications

The translational relevance of our findings is strengthened by the use of human-derived recombinant antibodies isolated from CSF samples of encephalitis patients (Kreye, Wenke et al. 2016). Exposure to human NMDAR-Ab was found to affect dendritic branching and maturation in cell cultures of rat cortical neurons (Okamoto, Takaki et al. 2022). A phenotypic assessment in a very comparable maternofetal transfer of human NMDAR-Ab resulted in altered nest building, poor motor coordination and impaired memory, that can resolve in adulthood (Garcia-Serra, Radosevic et al. 2021). Consequently, while the presence of the maladaptive brain state found in this study six weeks after exposure does not necessarily suggests a lasting and permanent shift in brain state, it might constitute a window of vulnerability, increasing the likelihood of a manifestation of a clinical phenotype upon exposure of a second, subsequent insult, e.g. by an environmental stressors or drug abuse. Consequently, we provide evidence for the presence of a maladaptive brain state which might contribute to or exacerbate the etiology of schizophrenia and other neurodevelopmental disorders.

## Acknowledgements

Panel A of figure 1 and the graphical abstract were created with BioRender.com.

## Author contributions

Conceptualization: SA, JD, HP, AS

Methodology: SA, JD, RGB, HB, JKu, JKr, CF

Investigation: SA, JD

Formal analysis: SA, HB, JKu, CF

Visualization: SA

Funding acquisition: AS Supervision: HP, AS

Writing – original draft: SA, JD, HP, AS

Writing – review & editing: SA, JD, HP, AS

## Funding

This study is supported by the Boehringer Ingelheim Foundation, the Leibniz Association and the German Research Foundation.

## Conflict of interest

The authors report no competing interests.

## Figures

**Fig. 6:**
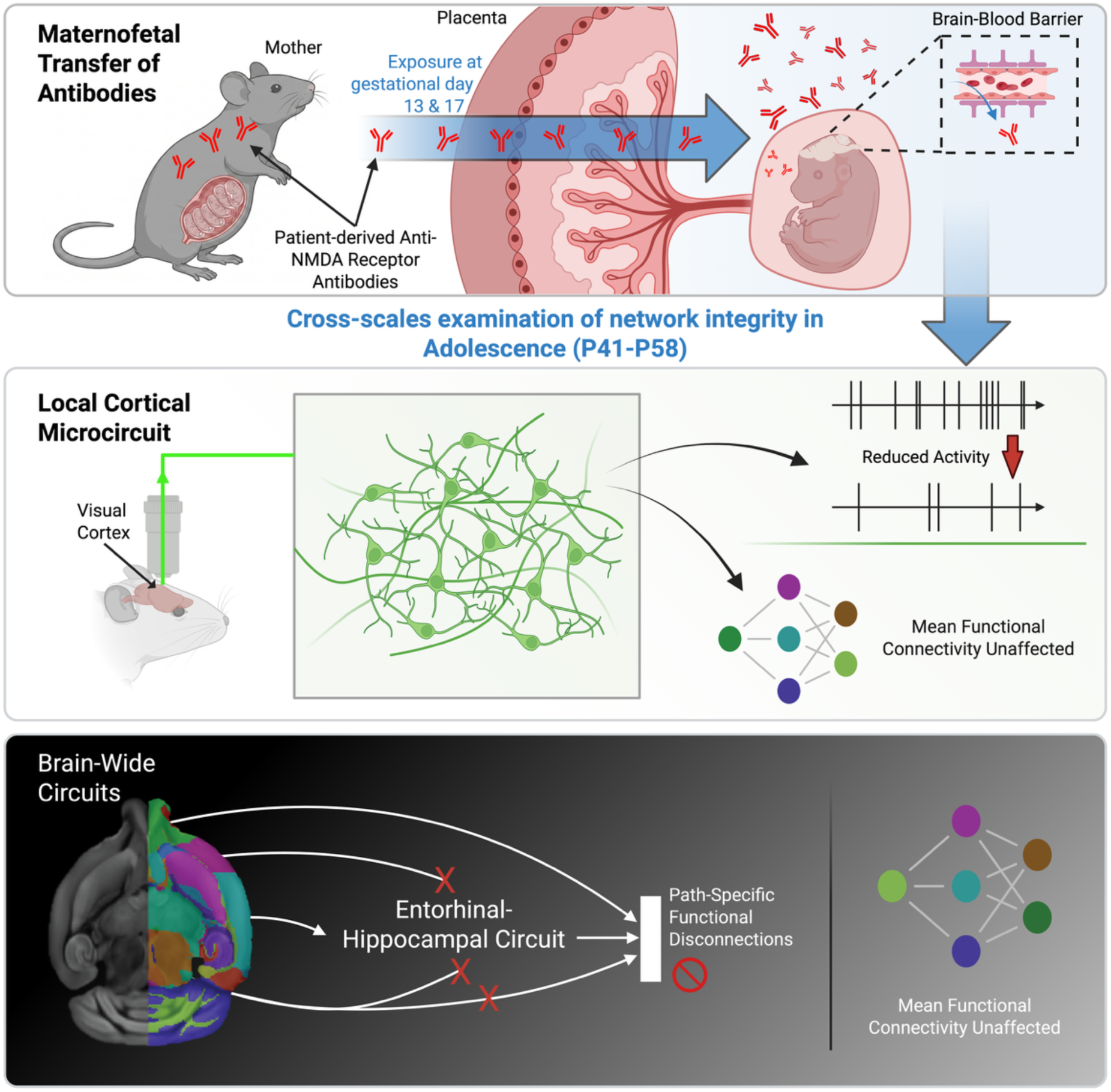
GRAPHICAL ABSTRACT.

**Supplementary Figure 1:**
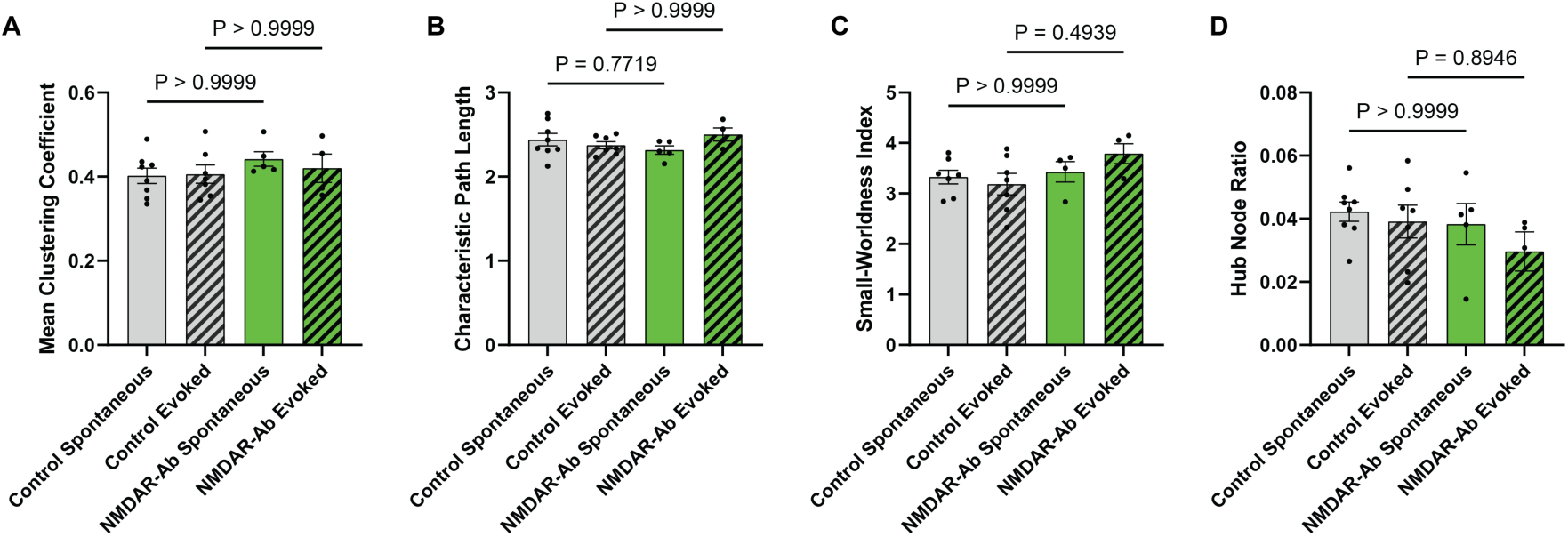
Graph Theoretical Analysis of two-photon recordings. **(A)** Clustering Coefficient remains stable under spontaneous (p>0.999) and evoked states (p>0.999, Dunn’s test). **(B)** Characteristic Path Length shows similar values between groups under spontaneous (p=0.772) and evoked states (p>0.999, Dunn’s test). **(C)** Small-Worldness Index, showing stable values under spontaneous (p>0.999) and evoked states (p=0.664, Dunn’s test). **(D)** Hub Neuron Ratio shows no significant differences under spontaneous (p>0.999) or evoked conditions (p=0.895, Dunn’s test).

**Supplementary Figure 2:**
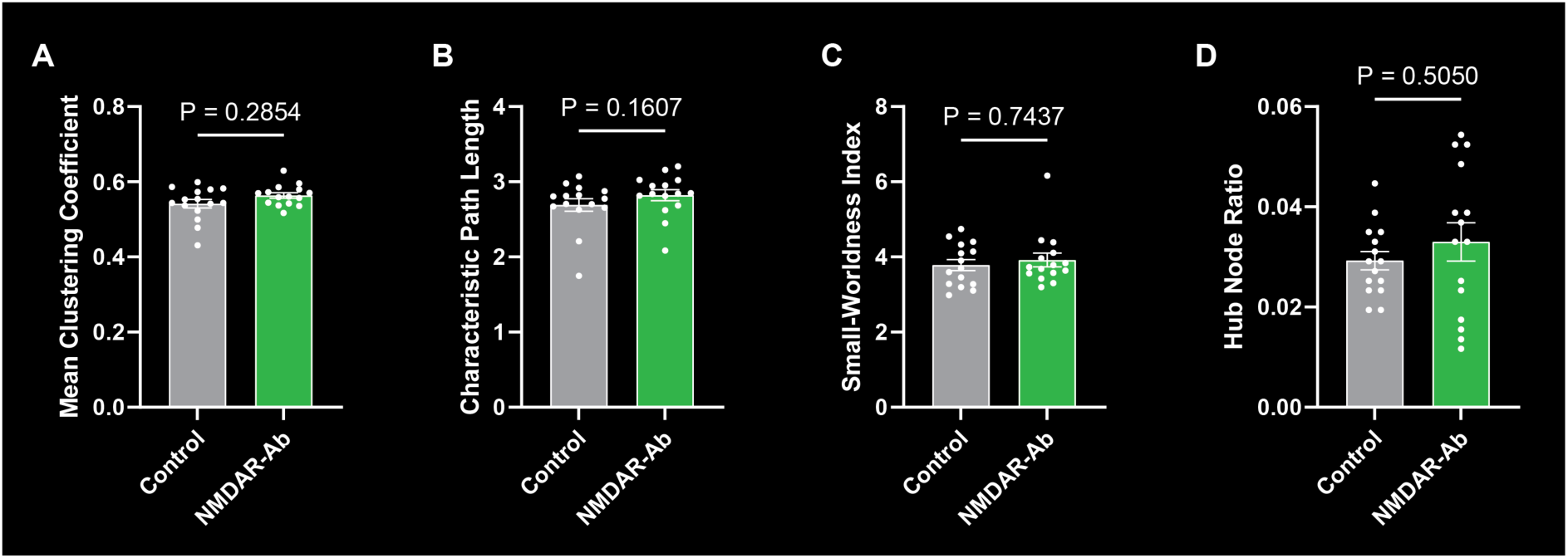
Graph Theoretical Analysis of fMRI recordings. **(A)** Clustering Coefficient is similar between groups (median: 0.5603 vs. 0.5439 in Control; Mann-Whitney test, p=0.285). **(B)** Characteristic Path Length shows no significant differences (median: 2.842 vs. 2.775 in Control; Mann-Whitney test, p=0.161). **(C)** Small-Worldness Index is stable between groups (median: 3.728 vs. 3.699 in Control; Mann-Whitney test, p=0.744). **(D)** Hub Region Ratio is unaffected by NMDAR-Ab exposure (median: 0.03301 vs. 0.02913 in Control; Mann-Whitney test, p=0.505).

## Notes

### Competing Interest Statement

The authors have declared no competing interest.

### Summary of Updates

New experimental data and results. We expanded the analyses of the two-photon calcium imaging and fMRI datasets. All figures have been revised to highlight the new findings.

